# Long-term decreased cannabinoid type-1 receptor activity restores specific neurological phenotypes in the Ts65Dn mouse model of Down syndrome

**DOI:** 10.1101/2021.11.22.469296

**Authors:** Anna Vázquez-Oliver, Silvia Pérez-García, Nieves Pizarro, Laura Molina-Porcel, Rafael de la Torre, Rafael Maldonado, Andrés Ozaita

**Affiliations:** Laboratory of Neuropharmacology-NeuroPhar. Department of Experimental and Health Sciences. University Pompeu Fabra, 08003 Barcelona, Spain; Integrative Pharmacology and Systems Neuroscience Research Group, Hospital del Mar Medical Research Institute, 08003 Barcelona, Spain; Alzheimer’s disease and other cognitive disorders unit. Neurology Service, Hospital Clínic, Spain; Neurological Tissue Bank, Biobanc-Hospital Clínic-IDIBAPS, Barcelona, Spain

**Author notes:** Corresponding author: Andrés Ozaita. Laboratory of Neuropharmacology-NeuroPhar. Department of Experimental and Health Sciences. Univ. Pompeu Fabra. C/Dr Aiguader 88, 08003 Barcelona, Spain. **Funding and disclosure** A.V.-O. is the recipient of a predoctoral fellowship from Jérôme Lejeune Foundation, France (Spanish Delegation); Bright Focus foundation (CA2018010) to L.M.; 2017 SGR 138 from the Departament d’Economia i Coneixement de la Generalitat de Catalunya (Spain) to R.T.; Plan Estatal RTI2018-099282-B-I00 (Spanish Ministerio de Ciencia, Innovación y Universidades) to A.O; The Jérôme Lejeune Foundation Grant#1896 (Cycle 2019B) to A.O; “Ministerio de Ciencia, Innovación y Universidades” (PID2020- 120029GB-I00/MICIN/AEI/10.13039/501100011033), “Ministerio de Sanidad, Servicios Sociales e Igualdad” (#RD16/0017/0020 to RM) and “Generalitat de Catalunya” (#2017 SGR-669 & #ICREA-Acadèmia 2015) to R.M.

**Keywords:** Down syndrome, CB1 cannabinoid receptor, rimonabant, Ts65Dn, memory

## Abstract

Intellectual disability is the most prevalent and limiting hallmark of Down syndrome (DS), without any pharmacological treatment available. Neurodegeneration and neuroinflammation are relevant neurological features of DS reaching to early development of Alzheimer’s disease. Preclinical evidence suggests that the endocannabinoid system, an important neuromodulator on cognition and neuroinflammation, could act as beneficial target in DS. Indeed, cannabinoid type-1 receptor (CB1R) activity was enhanced in the hippocampus of young-adult trisomic Ts65Dn mice, a well-characterized surrogate model of DS. In previous studies, inhibition of CB1R, was able to restore key neurological deficits in this mouse model. To determine the possible clinical relevance of this target, it is mandatory to evaluate the long-term consequences of attenuated CB1R activity and to minimize the possible side-effects associated to this mechanism. We found that CB1R expression was significantly enhanced in the hippocampus brains of aged DS subjects. Similarly, middle-aged trisomic mice showed enhanced CB1R expression. Long-term oral administration of a low dose of the CB1R specific antagonist rimonabant was administered to male and female Ts65Dn trisomic and wild-type mice from the time of weaning to 10 months, an age when signs of neurodegeneration have been described in the model. CB1R inhibition resulted in significant cognitive improvement in novel object-recognition memory in trisomic male and female mice, reaching a similar performance to that of wild-type littermates. Interestingly, this long-term rimonabant treatment modify locomotor activity, anxiety-like behavior, body weight or survival rates. Brain analysis at 10 months of age revealed noradrenergic and cholinergic neurodegeneration signs in trisomic mice that were not modified by the treatment, although the alterations in hippocampal microglia morphology shown by vehicle-treated trisomic mice was normalized in trisomic mice exposed to rimonabant. Altogether, our results demonstrate a sustained pro-cognitive effect of CB1R inhibition at doses that do not produce major side effects that could be associated to an anti-inflammatory action, suggesting a potential interest in this target of to preserve cognitive functionality in DS.

## Introduction

Down syndrome (DS) is the most significant genetic cause of intellectual disability affecting around 1 over 700-1,000 live births worldwide (de Graaf *et al*., 2021; de Graaf *et al*., 2020) and is produced by trisomy of human chromosome 21 (HSA21) (Lejeune *et al*., 1959). Intellectual disability is the most prevalent feature of DS an important limitation for people with DS independence especially in hippocampus-related cognitive domains (Dierssen, 2012; Pennington et al., 2003). The life expectancy for people with DS has markedly increased because of medical advances reaching to a median age at death of almost 60 years (De Graaf *et al*., 2017; Bayen *et al*., 2018). As a result, nowadays DS population is more prone to suffer from age-related comorbidities throughout their lives, such as an early-onset Alzheimer’s disease which is present in almost all adults with DS by age 65 (Mccarron *et al*., 2014). At the cellular level, aging subjects with DS show decreases in the density of noradrenergic neurons in the *locus coeruleus*, the main noradrenergic center in the brain (Mukhin *et al*., 2017) and in cholinergic neurons in the basal forebrain (Yates *et al*., 1983; Mann *et al*., 1985) as well as neuroinflammation (Wilcock and Griffin, 2013), all associated to the development of the Alzheimer’s disease pathology (Fortea *et al*., 2021).

Several animal models for DS have been established as a tool for investigating this condition based on the fact that HSA21 is orthologous to three distinct regions in murine chromosomes 10, 16, and 17 (Herault *et al*., 2017; Antonarakis *et al*., 2020). The Ts65Dn mouse model is the most-used mouse model for DS and it consists in a partial trisomy of murine chromosome 16 (from *App* to *Mx1*), covering more than 50% of genes in the homologous region of HSA21 (Davisson *et al*., 1993; Reeves *et al*., 1995). Importantly, Ts65Dn mice at 10 months of age also display common features with Alzheimer’s disease. Indeed, Ts65Dn mice show increased amyloid precursor protein (APP) expression, and degeneration of *locus coeruleus* noradrenergic neurons and basal forebrain cholinergic neurons, among others (Hamlett *et al*., 2015).

The endocannabinoid system (ECS) is a neuromodulatory system involved in synaptic homeostasis relevant for neuroinflammation and memory functions (Lutz, 2020). The ECS is composed by the cannabinoid receptors, mainly the cannabinoid type-1 and type-2 receptors (CB1R and CB2R, respectively), their endogenous ligands known as endocannabinoids, and the enzymes involved in their synthesis and degradation (Zou and Kumar, 2018). In the Ts65Dn model, previous studies showed that CB1R expression and function was enhanced in the hippocampus of young-adult Ts65Dn mice. Moreover, pharmacological and genetic CB1R inhibition restored memory deficits, synaptic plasticity, and adult neurogenesis in this mouse model at a young age (Navarro-Romero *et al*., 2019), although similar results have also been demonstrated after protecting 2-AG from enzymatic degradation (Lysenko *et al*., 2014). However, the possible efficacy of long-term treatments with CB1R antagonists in behavioral and in specific neurological phenotypes characteristic of aged Ts65Dn trisomic mice.

In this study, we found that CB1R expression is enhanced in hippocampal *post-mortem* samples of human subjects with DS further confirming CB1R as a potential therapeutic target. In addition, long-term pharmacological inhibition of CB1R with a low dose of rimonabant, enhanced memory in Ts65Dn mice at an age when this mouse model has a noticeable neurodegenerative and neuroinflammatory phenotype, the latter being sensitive to CB1R inhibition.

## Materials and methods

### Animals

All animal procedures were conducted following ARRIVE (Animals in Research: Reporting In Vivo Experiments) guidelines (Kilkenny *et al*., 2010) and standard ethical guidelines (European Communities Directive 2010/63/EU). Procedures were approved by the local ethical committee (Comité Ètic d’Experimentació Animal-Parc de Recerca Biomèdica de Barcelona, CEEA-PRBB) and local authorities (Generalitat de Catalunya). All experimental mice were bred at the Barcelona Biomedical Research Park (PRBB) Animal Facility. Ts65Dn experimental mice were obtained by repeated backcrossing Ts65Dn females to C57BL/6JEiJ x C3Sn.BLiA *Pde6b+*/DnJ F1 hybrid males. The parental generation was purchased from The Jackson Laboratory. Euploid littermates of Ts65Dn mice served as wild-type (WT) controls.

Mice were housed in Plexiglas cages with a maximum of 4 males or 5 female mice per cage in a temperature-controlled (21°C ± 1°C) (mean ± range) and humidity-controlled (55% ± 10%) environment. Lighting was maintained at 12 h cycles (light on at 8 AM; light off at 8 PM). All the experiments were conducted during the light phase in an experimental room at the animal facility. Food and water were available *ad libitum*. All behavioral experiments were performed by an observer blind to the genotype and treatment.

### Drug treatments

Rimonabant was purchased from Axon Medchem. Rimonabant was first prepared as a 40 mM stock solution in ethanol. Then, rimonabant stock solution was diluted to a final concentration of 10.8 μM in 0.3 % 2-hydroxypropyl-β-cyclodextrin in water. The compound was administered through the drinking bottle, and the same solution without rimonabant was used in littermate mice as control/placebo/vehicle condition. Animals with different treatment (vehicle or rimonabant) were maintained in different home-cages, while home-cages could hold both genotypes (Ts65Dn or WT). Mice were included in the study at the age of weaning (postnatal day 21, PND21) and randomly associated to one of the treatments. During the first 2.5 months of treatment mice received rimonabant 0.1 mg/kg/day. This dose was increased to the final concentration of 0.5 mg/kg/day by 3.5 months of age. The dosage was then maintained until mice were euthanized at 10 months of age.

### Behavioral tests

All behavioral tests were performed in a sound-attenuated room with dim illumination. A digital camera on top of the maze was used to record the sessions.

<u>The novel object-recognition memory test</u> was performed following a previously described protocol (Puighermanal *et al*., 2009). Briefly, novel object-recognition memory was assessed in a V-shape maze with dim illumination (3-5 lux). This task consists in 3 different phases (habituation, familiarization/training, and test) performed on 3 consecutive days for 9 min. On day 1, mice were habituated to the empty V-maze. Next day, mice were introduced in the V-maze where 2 identical objects were presented in the familiarization/training phase. Finally, the test was performed 24 h later, where 1 of the familiar objects was replaced for a novel object and the exploration time for both objects was recorded. Object exploration was defined as orientation of the nose toward the object at a distance < 2 cm. A discrimination index (DI) was calculated as the difference between the time spent exploring either the novel (Tn) or familiar (Tf) object divided by the total time spent exploring both objects: DI = (Tn − Tf) / (Tn + Tf).

<u>Locomotor activity</u> was assessed for 120 min. Individual locomotor activity boxes (9 × 20 × 11 cm) (Imetronic) were used in a low luminosity environment (5 lux). The total activity was detected by infrared sensors.

The <u>elevated plus maze test</u> was performed in a black Plexiglas apparatus with 4 arms (29 cm long x 5 cm wide), 2 open and 2 closed, set in cross from a neutral central square (5 cm x 5 cm) elevated 30 cm above the floor and indirectly illuminated from the top (40–50 lux in the open arms/4–6 lux in the close arms). Five-minute test sessions were performed, and total number of entries and the percentage of time spent in the open arms were used as a measure of anxiety-like behavior.

### Preparation of histological brain samples

Mice were deeply anesthetized by intraperitoneal injection (0.2 mL per 10 g body weight) of a mixture of ketamine (100 mg/kg) and xylazine (20 mg/kg). Then, mice were perfused intracardially with 4 % paraformaldehyde (PFA) in a 0.1M Na_2_HPO_4_/NaH_2_PO_4_ (pH 7.5) phosphate buffer (PB) using a peristaltic pump. The brains were removed from the skull and postfixed overnight at 4 °C in the same fixative solution. The next day, brain sections were moved to 30 % sucrose in PB solution. Brain sections (30 μm) were obtained with a sliding microtome and kept in a 5 % sucrose in PB solution at 4 °C until they were used for immunodetection.

### Immunofluorescence

Firstly, free-floating sections were washed three times (5 min each) in PB. Afterwards, the tissue was incubated in blocking buffer for 2 h which was made of 0.3 % Triton X-100 and 3 % normal donkey or goat serum diluted in 0.1 M PB, for neurodegeneration assessment and inflammation assessment respectively. Then, sections were incubated with one of the primary antibodies (mouse anti-TH (1:1,000, Sigma); rabbit anti-p75NTR (1:1,000, Millipore); mouse anti-Iba1 (1:500, Wako)) for 24 h at 4 °C. Then, sections were washed 3 times (10 min each) and subsequently incubated with the corresponding secondary fluorescent antibodies for 2 h at room temperature. As secondary antibodies were used goat anti-mouse Alexa Fluor 488 (1:1,000), donkey anti-rabbit Alexa Fluor 555 (1:500) and donkey anti-mouse Alexa Fluor 555 (1:500). Finally, the sections were washed 3 times (10 min each) and were mounted onto gelatin-coated slides with Fluoromont/DAPI mounting medium.

### Image acquisition and analysis

For TH+ cell counting, 4 coronal sections of the *locus coeruleus* were selected (from 5.34 to 5.52 posterior to Bregma) (Gould *et al*., 2012). Images of stained sections were obtained with a confocal microscope TCS SP8 LEICA (Leica Biosystems) using a dry objective (20× objective, 0.75 zoom) with a sequential line scan at 1024 × 1024-pixel resolution. The number of TH+ cells was manually quantified using Fiji software (Image J). The number of positive cells was calculated as the mean of total number of cells counted referred to the area of the *locus coeruleus* (μm^2^).

For p75NTR+ cell counting, systematic series of coronal sections (1 every 6 sections) per animal were selected, covering the rostral to caudal extension of the medial septum (from 1.18 and 0.38mm posterior to Bregma) (Gould *et al*., 2012).

The Leica DM6000B microscope was used (10x objective). To obtain the macro used for the quantification of p75NTR+ cells, firstly the Yen threshold was applied. Then an erosion and dilatation operation were applied with the command OPEN followed by remove outliers command (radius=2 and threshold=50). Finally, watershed command was executed and cells higher than 70μm^2^ were counted as positive. The number of positive cells was calculated as the mean of total number of cells counted referred to the area of the medial septum (μm^2^).

Iba1 staining was used to evaluate the microglial morphology. Confocal microscopy images of whole Z-stack from the slice were acquired in the *stratum radiatum*. Images of stained sections were obtained with a confocal microscope TCS SP5 STED LEICA (Leica Biosystems) using an immersion-oil objective (40× objective, 1.5 zoom) with a sequential line scan at 1024 × 1024-pixel resolution and with 0.3 μm depth intervals. The perimeter of the microglial soma was quantified in 20 cells per animal using Fiji software (Image J).

### Human samples slice histology, immunofluorescence, image acquisition and analysis

Brain samples (n=5 from subjects with DS, median age 64 and n=5 from typically developing subjects median age 66, all male) were obtained from the Neurological Tissue Bank, Biobanc-Hospital Clínic-IDIBAPS, Barcelona, Spain (Supplementary Table 1). Neuropathologic examination was performed according to standardized protocols (Borrego-Écija *et al*., 2021). Half-brain was fixed in formaldehyde solution for 3 weeks. Middle-posterior hippocampal region was embedded in paraffin and cut at 4 μm. The sections were placed in a BOND-MAX Automated Immunohistochemistry Stainer (Leica Biosystems Melbourne Pty Ltd, Melbourne, Australia). Tissues were deparaffinized and pre-treated with the Epitope Retrieval Solution 2 (EDTA-buffer pH8.8) at 98°C for 40 min. CB1R primary antibody (rabbit, 1:1,000, Immunogenes) was incubated for 60 min. Subsequently, tissues were incubated with polymer-HRP for 8 min and developed with DAB-Chromogen for 10 min. Slides were counterstained with hematoxylin.

Hippocampal subregions were defined using QuPath software (Bankhead *et al*., 2017) and following Allen Brain Atlas guidelines. Mean gray value optical density and % of occupied area were analyzed for CB1R and Nissl staining respectively, using Fiji software (Image J). The % of Nissl occupied area was quantified after the application of an optimal automatic threshold (Triangle dark) from Fiji software (ImageJ). Then, an erosion and dilatation operation were applied with OPEN command to erase unspecific signal. Ratio between CB1R optical density values and % of Nissl occupied area were expressed as a percentage of control group.

### Western Blotting

Mouse brain tissue was rapidly dissected, immediately frozen on dry ice and stored at −80 °C until used. Samples were processed following a protocol previously described (Ozaita *et al*., 2007) to obtain the total solubilized fraction and separated on SDS-PAGE gels and transferred into nitrocellulose membranes as previously described (Ozaita et al., 2007). The primary antibodies used were rabbit anti-CB1R (1:1,000, Immunogenes) and mouse anti-actin (1:50,000, MAB1501, Merck Millipore). Primary antibodies were detected with horseradish peroxidase conjugated anti-rabbit and anti-mouse antibodies and visualized by enhanced chemiluminescence detection (Luminata Forte Western HRP substrate, Merck Millipore). Digital images were acquired on a ChemiDoc XRS System (Bio-Rad) and quantified using The Quantity One software v4.6.3 (Bio-Rad). Optical density values for CB1R were normalized to actin optical density values as loading control in the same sample and expressed as a percentage of control group (WT).

### Rimonabant detection

For the analysis of mice brain content, tissues (40-80 mg) where weighted and homogenized with 1 mL grinder dounce (Wheaton, USA) in two steps: first by adding 400 μL of HCOOH 0.1 %, (thirty movements “loose”, followed by thirty movements “tight” were used for homogenization) followed by protein precipitation with 800 μL of ethanol. The mixture was centrifuged 10 min at 15,700 g, 4 °C, and supernatant was recovered and stored at −20 °C until use. Then, 100 μL of supernatant was mixed with 25 μL of the internal standard (rimonabant-d10, 0.004 μg/mL in ethanol, Bertin technologies, Montigny le Bretonneux, France) and 4 μL of the final mixture were used for the HPLC-MS/MS analysis.

Samples were analyzed in an Acquity UPLC System (Waters Associates, Milford, MA, USA) coupled to a mass spectrometer (Quattro Premier, Watters Associates). Chromatographic separation was carried out in an Acquity BEH C18 (100 mm x 2.1 mm i.d., 1.7 μm) (Waters Associates) at a flow rate of 0.4 mL/min. Ammonium formate (1 mM)-HCOOH 0.01% (A) and MeOH with ammonium formate (1 mM)-HCOOH 0.01%(B) were used as mobile phases. After keeping 40%B for 0.5 min, the gradient was increased to 95%B in 3 min and maintained at 95%B for 1 minute after going back to initial conditions. Detection of analytes was done by the selected reaction monitoring (SRM) method, being the transitions used for identification and quantification (in bold) for each compound as follows: 465→84, 99, 365 (rimonabant); 475→94, 365 (rimonabant-d10).

### Experimental design and statistical analysis

Sample size choice was based on previous studies with similar experimental approached (Busquets-Garcia *et al*., 2013, 2016) and it is indicated in figure legends for each experiment. Data were analyzed with GraphPad Software using unpaired Student’s t-test or two-way ANOVA for multiple group comparisons. Subsequent *post-hoc* analysis (Bonferroni) was used when significance in interaction between factors was found. The Pearson correlation coefficient was used to analyze the relationship between discrimination index and the area of microglial soma. Comparisons were considered statistically significant when p < .05. Outliers (±2 s.d. from the mean) were excluded.

## Results

### CB1R expression is enhanced in hippocampal tissue of subjects with DS and middle aged trisomic Ts65Dn

Previous studies in our laboratory described that the expression of CB1R was increased in the hippocampus of young-adult Ts65Dn male mice (Navarro-Romero *et al*., 2019). We hypothesized that hippocampal CB1R expression could be also enhanced in human subjects with DS. Therefore, we immunodetected and quantified the expression of CB1R in brain slices corresponding to the hippocampal formation of aged subjects with DS and sex, age, and *post-mortem* interval-matched controls (Supplementary Table 1). CB1R immunodetection was non-significantly increased when all subregions of the hippocampus were considered together (Figure 1A). Interestingly, the analysis of the CB1R immunoreactivity in the dentate gyrus showed a significant increase in samples from DS subjects (Figure 1B). Protein expression analysis of CB1R in hippocampus of trisomic Ts65Dn middle-aged mice also showed a significant increase in comparison to WT litter mates’ mice (Figure 1C). Together, these data revealed that CB1R overexpression is shared in both human subjects with DS and middle-aged trisomic Ts65Dn mice.

**Figure 1.**
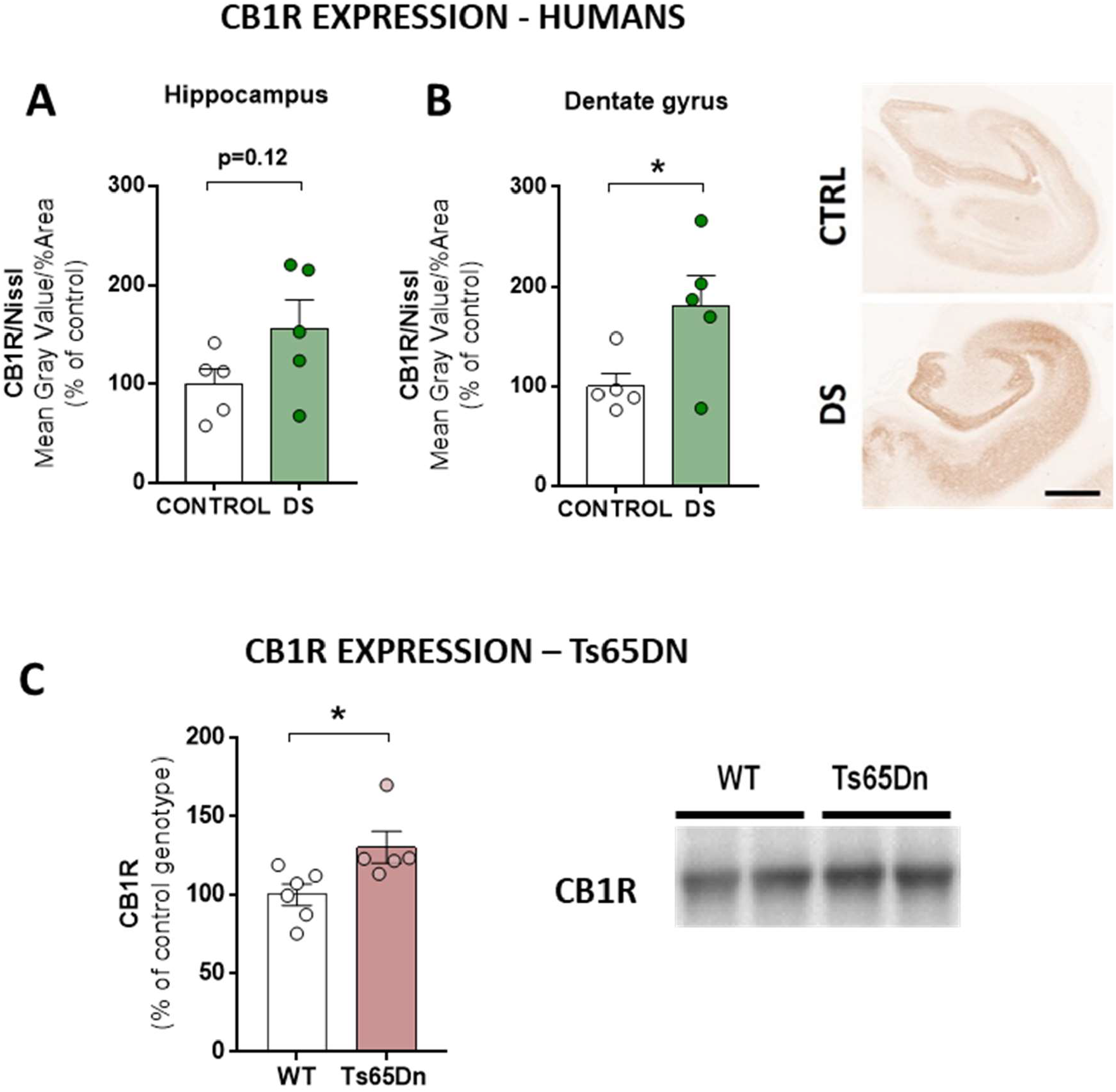
CB1R expression is enhanced in human *post-mortem* samples of aged Down syndrome subjects and in hippocampus of middle-aged Ts65Dn mice. (A-B) Representative images and average intensity of CB1R expression in the hippocampus (A) and dentate gyrus (B) of Control and subjects with DS (Control, n=5, DS=5) (scale bar = 2 mm). (C) Representative immunoblots and quantification of CB1R in hippocampus from WT and Ts65Dn mice of 10 months of age (WT, n=6, Ts65Dn, n=4). Distribution of individual data with mean ± s.e.m. * *p* < .05 (genotype effect) by Student’s *t*-test.

### Sustained oral administration of a low dose of rimonabant improves memory performance in young-adult and middle-aged Ts65Dn mice

We then tested whether a long-term sustained pharmacological intervention with a low dose of the selective CB1R antagonist rimonabant (Rinaldi-Carmona *et al*., 1994), would be suitable to improve memory performance in Ts65Dn trisomic mice. To this aim, we first established the conditions to perform an oral treatment to deliver rimonabant in the drinking water. Rimonabant was stable in solution at room temperature for at least 4 days (data not shown), so bottle content was changed every 4 days for the length of the treatment. Both male and female mice (Ts65Dn and WT littermates) received the treatment (rimonabant or vehicle) through the drinking water from PND21 until 10 months of age, at this time brain samples were collected. During treatment, mice were analyzed in their behavioral response to assess different aspects of general activity as that could be of relevance for the possible side effects associated to CB1R blockade as well as for the efficacy of the treatment at the cognitive level (Figure 2A). First, mice were tested at 4 months of age using the novel object recognition (NOR) task. We observed that Ts65Dn male mice treated with vehicle presented a deficit in this task, as expected (Reeves *et al*., 1995; Fernandez *et al*., 2007). Instead, Ts65Dn male mice treated with rimonabant presented a better cognitive performance, comparable to controls treated with vehicle (Figure 2B). This observation confirmed that sustained inhibition of CB1R does not show tolerance to the mnemonic effects observed in trisomic mice.

**Figure 2.**
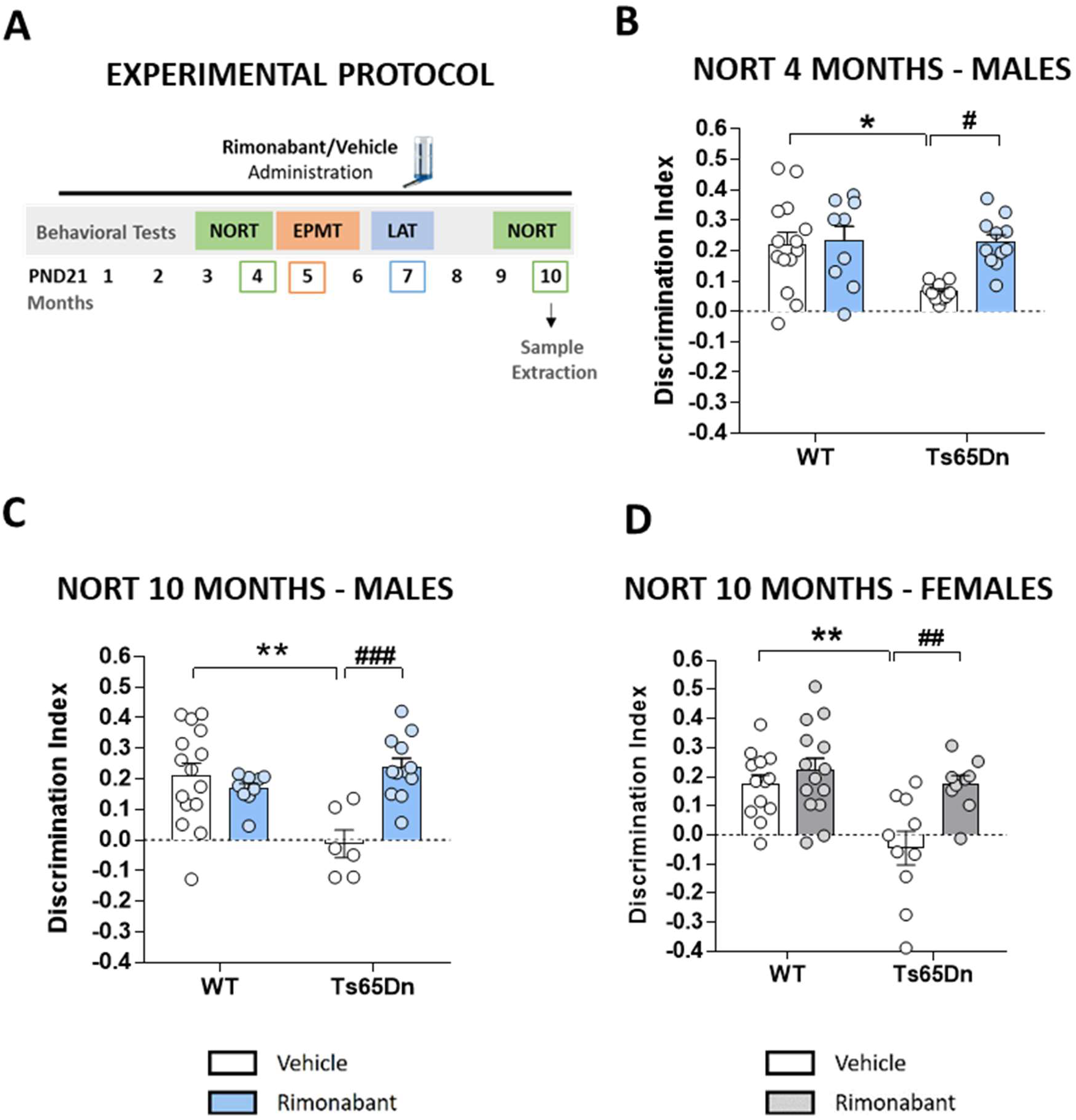
Long-term CB1R inhibition improves memory performance in young and in middle-aged male and female Ts65Dn mice. (A) Schematic representation of the experimental protocol. EPMT = elevated plus maze test, LAT = locomotor activity test, NORT = novel object recognition test, PND21 = postnatal day 21. (B) Discrimination index in novel object-recognition test (NORT) of WT and Ts65Dn male of 4 months of age with VEH or RIM (0.5 mg/kg/day) (WT VEH, n=14; WT RIM, n=9; Ts65Dn VEH n=10; Ts65Dn RIM n=11). (C-D) Discrimination index in novel object-recognition test (NORT) of WT and Ts65Dn male (C) and female (D) mice treated of 10 months of age with VEH or RIM (0.5 mg/kg/day) (males: WT VEH, n=15; WT RIM, n=9; Ts65Dn VEH n=6; Ts65Dn RIM n=12; females: WT VEH, n=13; WT RIM, n=14; Ts65Dn VEH n=10; Ts65Dn RIM n=9). Distribution of individual data with mean ± s.e.m. * *p* < .05, ** *p* < .01 (genotype effect); # *p* < .05, ## *p* < .01, ### *p* < .001 (treatment effect) by Bonferroni *post-hoc* test following two-way ANOVA.

To assess any possible behavioral disruption that could be related to potential side effects associated to CB1R blockade, we first evaluated anxiety-like behavior using the elevated plus maze task (EPMT) in the same cohort of mice. Ts65Dn trisomic male mice showed a low anxiety-like phenotype and rimonabant treatment did not modify such responses (supplementary Figure 1A). In contrast, rimonabant treatment increased anxiety-like behavior in WT female mice at this time point (supplementary Figure 1B). Interestingly, we observed a significant increase in the total number of entries in Ts65Dn trisomic mice compared to WT that was not altered by rimonabant treatment (supplementary Figure 1C and 1D). Locomotor activity (LAT) assessment of the same cohorts revealed an hyperlocomotor phenotype in Ts65Dn trisomic mice that was not modified by rimonabant exposure (supplementary Figure 2A and 2B). Together, these findings show that chronic long-term rimonabant therapy is effective in restoring NOR memory in young-adult Ts65Dn trisomic mice without compromising other behavioral features in these trisomic animals.

Because Ts65Dn mice display a noticeable neurodegenerative phenotype that resembles that of middle-aged subjects with DS, NOR memory was evaluated again in the same cohort of mice at 10 months of age. Interestingly, no tolerance was developed over the months of treatment since memory improvement was detected in the same cohort of rimonabant-treated male and female Ts65Dn mice of this second time point (Figure 2C and 2D). Remarkably, NOR test did not reveal an effect of the treatment in WT mice (Figure 2C and 2D) demonstrating that blocking CB1R specifically enhances hippocampal-dependent memory in Ts65Dn mice. Furthermore, the presence of rimonabant was detected in brain samples of treated mice at that time (mean ± s.e.m., Rimonabant treated groups = 12. 6 ± 6.0 pg/mg).

### Noradrenergic and cholinergic alterations are not prevented by long-term rimonabant treatment

Noradrenergic neurons in the *locus coeruleus* (LC-NE neurons) suffer degeneration in DS (Mukhin *et al*., 2017). We assessed whether long-term rimonabant treatment could alter LC-NE neurodegeneration in Ts65Dn trisomic mice where we revealed that this treatment improved memory performance. We used tyrosine hydroxylase (TH), the limiting enzyme in the synthesis of dopamine and norepinephrine (Vecchio *et al*., 2021), as a specific marker of LC-NE neurons. Analysis of LC-NE neurons revealed a significant decrease in the density of TH+ neurons in this region of Ts65Dn male mice, replicating similar results of previous studies (Salehi *et al*., 2009). Notably, LC-NE neuron loss was not prevented by long-term rimonabant treatment (Figure 3A).

**Figure 3.**
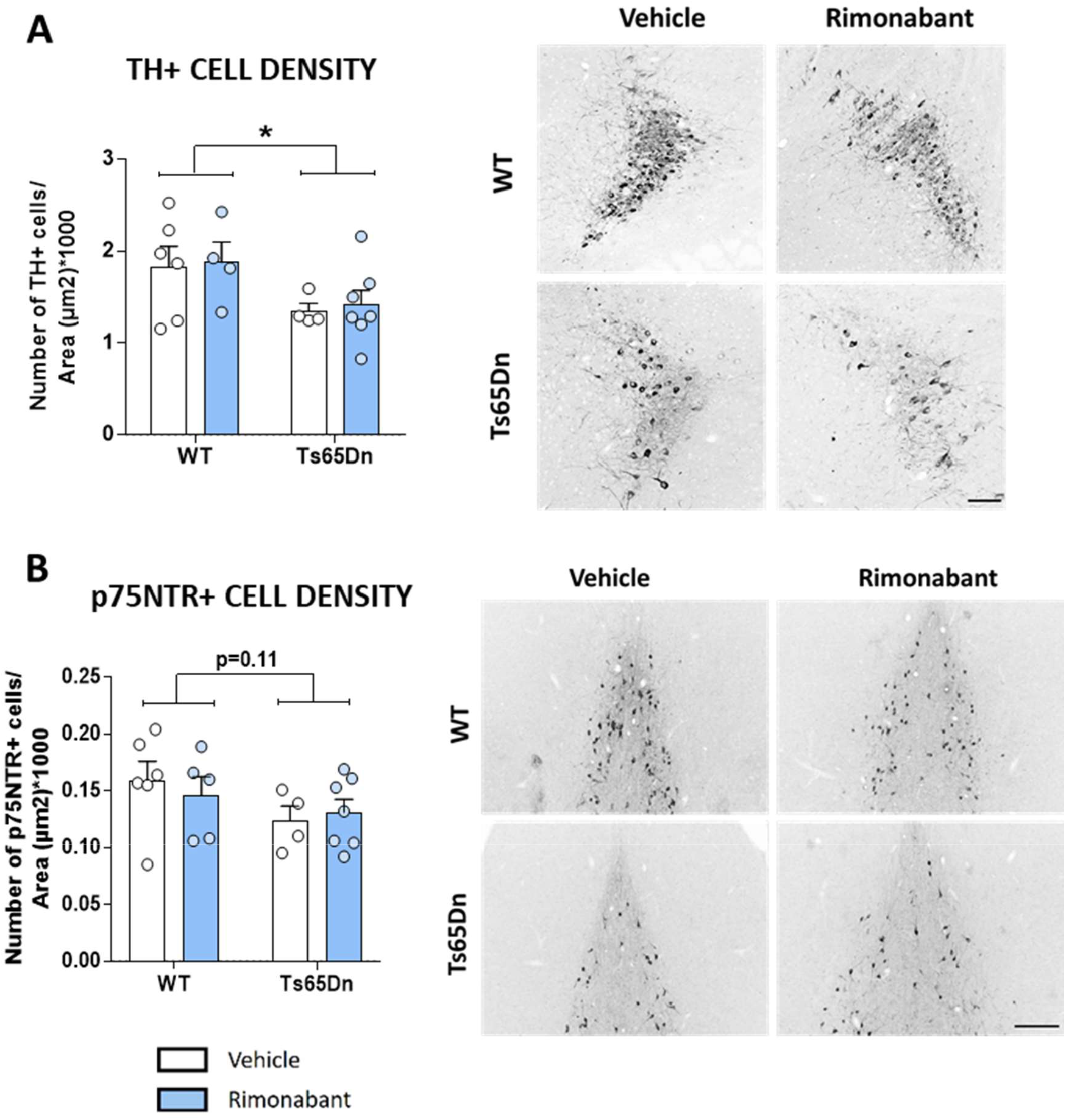
Long-term CB1R pharmacological inhibition did not modify the degeneration of cholinergic and adrenergic neurons in male Ts65Dn mice. (A) Representative grey scale confocal images and average density of TH+ cells in the *locus coeruleus* of WT and Ts65Dn mice of 10 months of age treated with VEH or RIM (0.5 mg/kg/day) (WT VEH, n=6; WT RIM, n=4; Ts65Dn VEH n=4; Ts65Dn RIM n=7) (Scale bar= 100 μm). (B) Representative grey scale images and average density of p75NTR+ cells in the medial septum of the basal forebrain of WT and Ts65Dn mice of 10 months of age with VEH or RIM (0.5 mg/kg/day) (WT VEH, n=6; WT RIM, n=5; Ts65Dn VEH n=4; Ts65Dn RIM n=7) (Scale bar= 200 μm). Distribution of individual data with mean ± s.e.m. * *p* < .05 (genotype effect) by two-way ANOVA.

Similarly, basal forebrain cholinergic neurons, the main source of acetylcholine to the hippocampus, undergo age-related degeneration in DS (Yates *et al*., 1983; Mann *et al*., 1985). Ts65Dn trisomic male mice show age-related atrophy and loss of cholinergic neurons particularly in the medial septum (Hamlett *et al*., 2015) as measured using immunostaining for either p75 neurotrophin receptor (p75NTR) or choline acetyltransferase (Salehi *et al*., 2006). We used p75NTR as a cholinergic marker. Under our experimental conditions, we found that Ts65Dn mice exhibit a non-significant tendency to decrease in the density of cholinergic cells in the medial septum of Ts65Dn compared to WT (Figure 3B), in line with previous studies (Granholm *et al*., 2000; Hunter *et al*., 2004b). Nevertheless, this trend to cholinergic degeneration was not modified by rimonabant treatment (Figure 3B). Together, these findings suggest that the effects of rimonabant treatment improving memory performance in aged Ts65Dn mice is independent from preventing the neurological changes revealed in noradrenergic and cholinergic neurons in these brain areas.

### Anomalous microglial morphology in Ts65Dn trisomic mice is normalized by long-term treatment with rimonabant

Together with early neurodegeneration, brain inflammation is a relevant neurological feature in Alzheimer’s disease in DS (Wilcock and Griffin, 2013; Wilcock *et al*., 2015). Microglial cells are the major mediators of the neuroinflammatory response in the brain and different studies have observed an increase in microglial reactivity in adult Ts65Dn mice (Hunter *et al*., 2004b; Lomoio *et al*., 2009; Illouz *et al*., 2019; Hamlett *et al*., 2020a) and in subjects with DS (Pinto *et al*., 2020). In the same previously studied cohort of mice, we observed an increase in microglial soma size, measured as cell body area, in the hippocampus of trisomic mice. This microglial activation was reversed to control values in long-term rimonabant-treated trisomic mice (Figure 4A). Moreover, microglia reactivity, negatively correlated with NOR memory performance (Figure 4B), pointing to a potential relationship between memory performance improvement and diminished microglial reactivity due to rimonabant treatment. Together, these results indicate that long-term CB1R pharmacological inhibition normalized microglia morphology in the hippocampus of trisomic mice.

**Figure 4.**
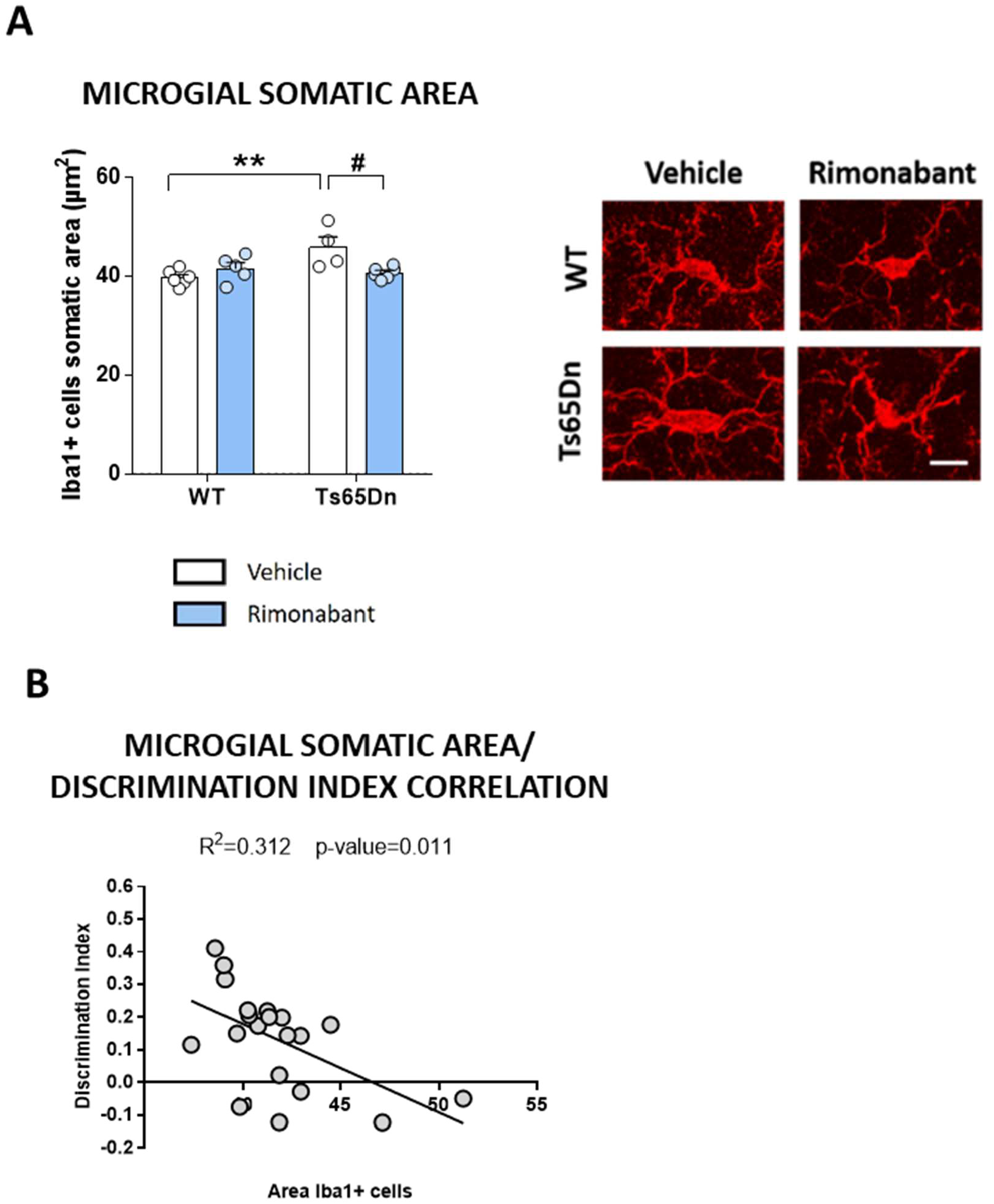
Long-term CB1R pharmacological inhibition reduced microglia reactivity in the hippocampus of male Ts65Dn mice. (A) Representative confocal images and average microglial somatic area of Iba1+ cells in the hippocampus of WT and Ts65Dn mice of 10 months of age with VEH or RIM (0.5 mg/kg/day) (WT VEH, n=6; WT RIM, n=5; Ts65Dn VEH n=4; Ts65Dn RIM n=6) (scale bar=10 μm). (B) Correlation between discrimination index in the NORT and microglial activation (area of the soma) in the hippocampus. Distribution of individual data with mean ± s.e.m. ** *p* < .01 (genotype effect); # *p* < .05 (treatment effect) by Bonferroni *post-hoc* test following two-way ANOVA.

## Discussion

Intellectual disability in DS is characterized by learning and memory impairments together with age-related comorbidities, such as Alzheimer’s disease. Currently, there is no effective treatment to circumvent the genetic alterations in DS. Therefore, there is an urgent need to investigate novel therapeutic targets. In this study, we describe, for the first time to the best of our knowledge, a dysregulation in the expression of CB1R in the hippocampus of human subjects with DS and Ts65Dn middle-aged trisomic mice, revealing CB1R as a promising target for long-term treatments to enhance hippocampal-dependent memory in this disease.

We previously described that CB1R expression and function were upregulated in young-adult Ts65Dn trisomic mice (Navarro-Romero *et al*., 2019), but whether this feature is observed in human subjects with DS had not been previously addressed. Therefore, we analyzed CB1R immunodetection in the hippocampus of aged subjects with DS compared to age-matched healthy controls. Interestingly, 4 of 5 subjects with DS involved in this study suffered from Alzheimer’s disease. A significant enhancement of CB1R immunodetection was found in the dentate gyrus of the hippocampus, whereas a non-significant increased was observed in the whole hippocampus. These findings further underlie the potential interest of diminishing CB1R function as a suitable pharmacological approach for the treatment of memory impairment in DS, as we previously suggested (Navarro-Romero *et al*., 2019). We also observed that hippocampal CB1R expression was enhanced in homogenates of middle-aged Ts65Dn trisomic mice. At this age, Ts65Dn trisomic mice show signs of a neurodegenerative phenotype (Hamlett *et al*., 2015). Studies of CB1R expression in the brains of Alzheimer’s disease patients have been inconsistent. Indeed, CB1R has been shown to be downregulated, upregulated, and unaltered in these patients (Westlake *et al*., 1994; Lee *et al*., 2010; Ahmad *et al*., 2014; Manuel *et al*., 2014). Therefore, the enhanced hippocampal CB1R expression could be an up-to-now undescribed novel feature in subjects with DS and supports the potential interest of therapeutical approaches to attenuate CB1R function. Further studies will be needed to determine whether CB1R overexpression appears in younger subjects with DS as a common feature of the DS condition or whether it is associated to the neurodegenerative status in humans.

We then assessed whether a long-term CB1R pharmacological inhibition intervention would be efficacious for the improvement of memory performance in the mouse model. Rimonabant, a well stablished CB1R specific antagonist (Rinaldi-Carmona *et al*., 1994), reached the market (Acomplia®, Sanofi Aventis) as a treatment for obesity since high doses of rimonabant were found to reduce food intake and food reinforcing properties (Pacher, 2006). However, Acomplia® was withdrawn from the market due to psychiatric side effects affecting a subpopulation of obese patients (Christensen *et al*., 2007; Rucker *et al*., 2007). At high dosages, rimonabant has inverse agonist properties on CB1R, which has been hypothesized to be the cause of some of the negative effects associated to the treatment with this CB1R antagonist (Landsman *et al*., 1997; Silvestri and Di Marzo, 2012). Therefore, the use of low doses of rimonabant is mandatory to evaluate any possible therapeutic interest of this target in order to obtain results with potential translational value.

We observed that long-term rimonabant exposure, using at a dose 20 times lower that that used to reveal the anti-obesity properties in diet-induced obesity (Martín-García *et al*., 2010), was well tolerated and did not modify the body weight in mice compared to vehicle-treated animal (data not shown). Notably, recognition memory performance was preserved in both male and female young-adult and middle-aged Ts65Dn trisomic treated mice discarding tolerance processes, which have been observed to be key adverse feature to reduce the effectiveness of other treatments envisioned to improve cognition in other intellectual disability disorders (Stoppel *et al*., 2021). Our results of long-term exposure further support the relevance of CB1R to tackle cognitive impairment in DS as observed in sub-chronic treatments previously assessed (Navarro-Romero *et al*., 2019), and open the possibility to evaluate this approach in other intellectual disability disorders where low doses of CB1R antagonists have also been found effective, such as in fragile X syndrome (Busquets-Garcia et al., 2013; Gomis-González et al., 2016).

Other studies have found that increasing the endocannabinoid 2-arachidonoil glycerol (2-AG), by inhibiting the metabolizing enzyme monoacylglycerol lipase (MAGL) through JZL184, has favorable benefits on cognitive function and synaptic plasticity in 11-month-old Ts65Dn trisomic mice (Lysenko *et al*., 2014). Although the approach used in our study with middle-aged Ts65Dn mice would appear to be contradictory, both rimonabant and JZL184 treatments could reduce CB1R functionality. Indeed, JZL184 treatment in mice, or mice knockout for MAGL, has been shown to cause CB1R desensitization has been reported due to the extra 2-AG available (Schlosburg *et al*., 2010; Bernal-Chico *et al*., 2015). Therefore, both therapies, through different mechanisms, may eventually reduce CB1R functionality after long term exposure. Furthermore, the possibility that JZL184 is acting through a CB1R-independent mechanism cannot be discarded.

Middle-aged human adults with DS present neuropathological alterations common to Alzheimer’s disease, involving the degeneration of noradrenergic neurons in the *locus coeruleus* and basal forebrain cholinergic neurons (Ballard *et al*., 2016; Fortea *et al*., 2021), features that are reproduced by middle-aged Ts65Dn mice (Granholm *et al*., 2000; Salehi *et al*., 2009; Hamlett *et al*., 2015). Therefore, we have analyzed the neurodegenerative features of Ts65Dn mice after long-term rimonabant exposure. Interestingly, we observed that Ts65Dn treated with rimonabant presented evident noradrenergic cell loss and an emerging cholinergic impairment to a similar extent that Ts65Dn treated with vehicle. These results discard an effect of rimonabant in these particular neuronal populations to explain its positive effects on cognition in Ts65Dn mice.

Previous research has revealed that endocannabinoids are dysregulated in Alzheimer’s disease and contribute to the disorder’s progression (Cristino *et al*., 2020). Because research shows conflicting results, it is difficult to determine if an increase or reduction in cannabinoid tone is associated to an improvement in pathology. For instance, CB1R and/or CB2R agonists improved memory and/or cognitive deficits in Tg2576 mice, APP/PS1 mice (Aso *et al*., 2013), and in rodents receiving intracerebral injections of Aβ (Ramírez *et al*., 2005; Martín-Moreno *et al*., 2011). Conversely, CB1R antagonism protected against Aβ-induced memory impairment in mice (Mazzola *et al*., 2003), suggesting that activation of CB1R by endocannabinoids inhibits neurotoxicity, but may worsen its long-term consequences (such as reduced acetylcholine signaling) that lead to cognitive impairment.

Given that long-term rimonabant treatment prevented memory deficits in the Ts65Dn mice, we explored other neurological parameters relevant for cognitive performance, such as neuroinflammation. Neuroinflammation is a major contributor to neurodegenerative disorders (Colonna and Butovsky, 2017; Yin *et al*., 2017) including Alzheimer’s disease in DS (Wilcock, 2012; Wilcock *et al*., 2015; Flores-Aguilar *et al*., 2020; Pinto *et al*., 2020). Interestingly, recent studies also demonstrated that a reduction in microglial reactivity was associated to improvements in learning and memory in a mouse model of DS (Pinto *et al*., 2020). We found that Ts65Dn trisomic mice displayed an increase in microglial soma size, which is associated to a reactive type of microglia (Helmut *et al*., 2011), in agreement with previous studies in microglial populations in the Ts65Dn model (Hunter *et al*., 2004b). Interestingly, rimonabant-treated Ts65Dn trisomic mice that showed regular recognition memory exhibited as well similar microglial body area that control mice. This anti-inflammatory effect of rimonabant could be directly impinged onto microglial cells or could be secondary to the effect of rimonabant on neuronal circuits that promote microglial reactivity. Notably, the microglial soma area correlated with memory performance in the novel-object recognition test. Previous studies have linked microglial activation and cholinergic cell loss (Hunter *et al*., 2004b), although we did not find an association between these two brain alterations after long-term rimonabant treatment in our study. One possibility would be that the neuroinflammatory effect is limited to the hippocampal region, with no effect in degenerative regions like the basal forebrain. As a result, these findings suggest a link between improved memory function and reduced microglial reactivity after rimonabant treatment.

At the present moment, there are no approved treatments for intellectual disability in DS in spite of the large amounts of preclinical studies that have been performed (Hamlett *et al*., 2015; Stagni *et al*., 2015). In our current study, we maximized the potential translational value of our experimental approach by taking numerous variables into consideration. First, we studied the status of CB1R in DS human hippocampus, and we found interesting parallelisms with the preclinical models. Second, we performed our study in the most used preclinical model for DS, the Ts65Dn, which predictive validity was recently proven for novel experimental approaches to enhancing memory performance in subjects with DS (de la Torre *et al*., 2014; de la Torre *et al*., 2016). In addition, Ts65Dn mouse model is the only model which neurodegenerative phenotype is widely described, necessary for the assessment of long-term effects of our treatment (Herault *et al*., 2017; Antonarakis *et al*., 2020). Third, we assessed rimonabant efficacy in male and female mice, presenting comparable sex results. Altogether, our results reinforce the potential interest of decreasing CB1R activity to maintain cognitive function and prevent specific neurological deficits in DS and expands our understanding of the potential use of CB1R inhibition for long-term periods in the treatment of this disease.

## Acknowledgments

We are grateful to Raquel Martín, Marta Linares, Dulce Real, Jolita Jancyte and Francisco Porrón for expert technical assistance. We are grateful to Lydia García-Serrano, Gabriela Bordeanu and the Laboratori de Neurofarmacologia-NeuroPhar for helpful discussion. We are indebted to the Biobanc-Hospital Clinic-Institut d’Investigacions Biomèdiques August Pi I Sunyer (IDIBAPS) for samples and data procurement.

