## Supplemental Material for "Long-term decreased cannabinoid type-1 receptor activity restores specific neurological phenotypes in the Ts65Dn mouse model of Down syndrome"

| Case Number | Diagnosis | Age (years) | Gender | Clinical diagnosis of dementia |
|---|---|---|---|---|
| 1 | Control | 66 | M | Absent |
| 2 | Control | 70 | M | Absent |
| 3 | Control | 64 | M | Absent |
| 4 | Control | 58 | M | Absent |
| 5 | Control | 76 | M | Absent |
| 6 | Down syndrome | 62 | M | Present |
| 7 | Down syndrome | 59 | M | Present |
| 8 | Down syndrome | 64 | M | Present |
| 9 | Down syndrome | 61 | M | Present |
| 10 | Down syndrome | 71 | M | Absent |

**Supplementary Table 1. Human samples used for immunohistochemistry for CB1R in Figures 1A-B.**

### ANXIETY-LIKE BEHAVIOR

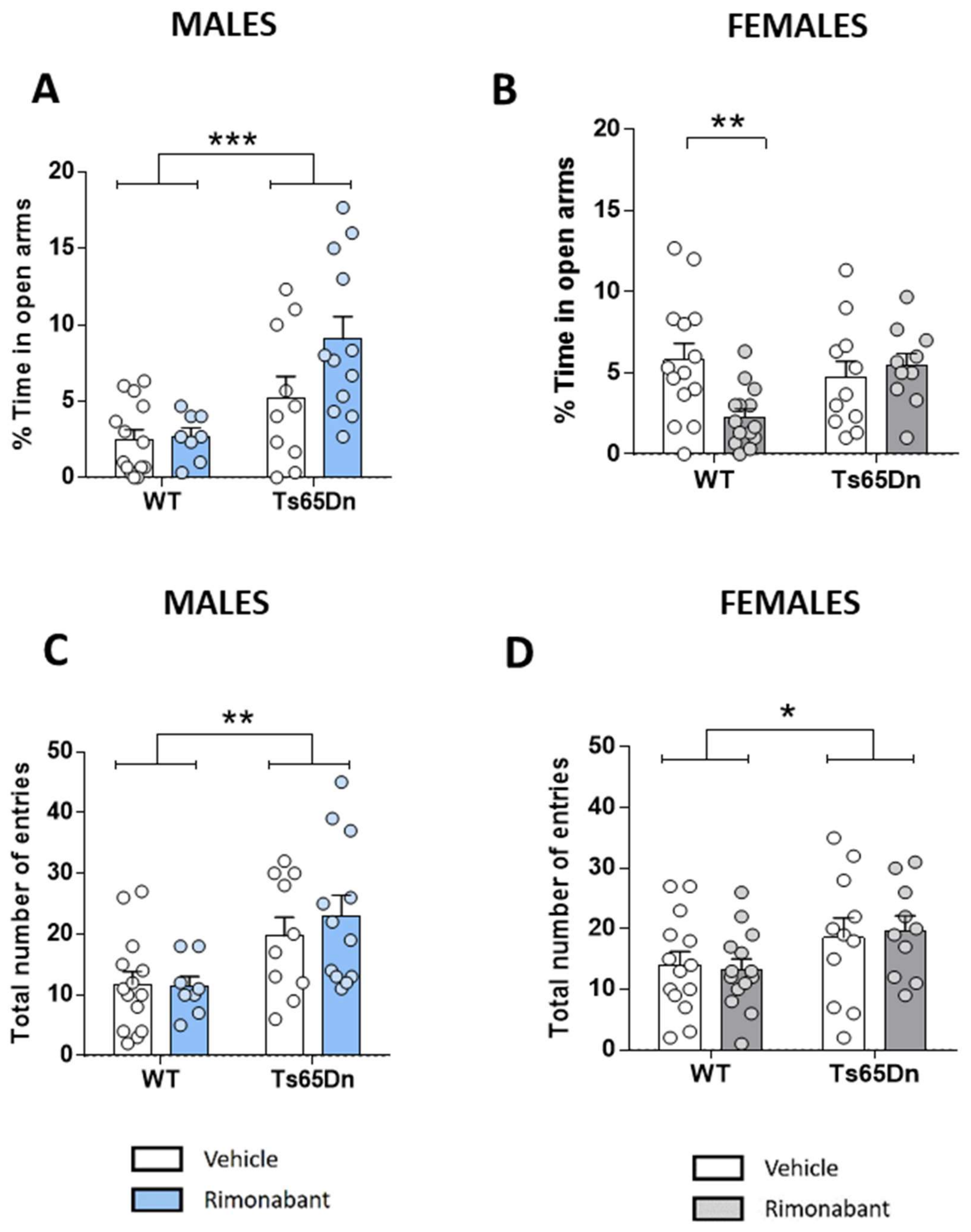

**Supplementary Figure 1. Long-term CB1R pharmacological inhibition does not modify anxiety-like behavior in Ts65Dn.** (A-B) Percentage of time in open arms and number of total entries in the elevated plus maze test of male (A) and female (B) WT and Ts65Dn mice of 7 months of age treated with VEH or RIM (0.5 mg/kg/day) (males: WT VEH, n=14; WT RIM, n=8; Ts65Dn VEH n=10; Ts65Dn RIM n=12; females: WT VEH, n=14; WT RIM, n=14; Ts65Dn VEH n=11; Ts65Dn RIM n=10). Distribution of individual data with mean ± s.e.m., *** $p < .001$, ** $p < .01$, * $p < .05$ (genotype effect) by Bonferroni *post-hoc* test following two-way ANOVA.

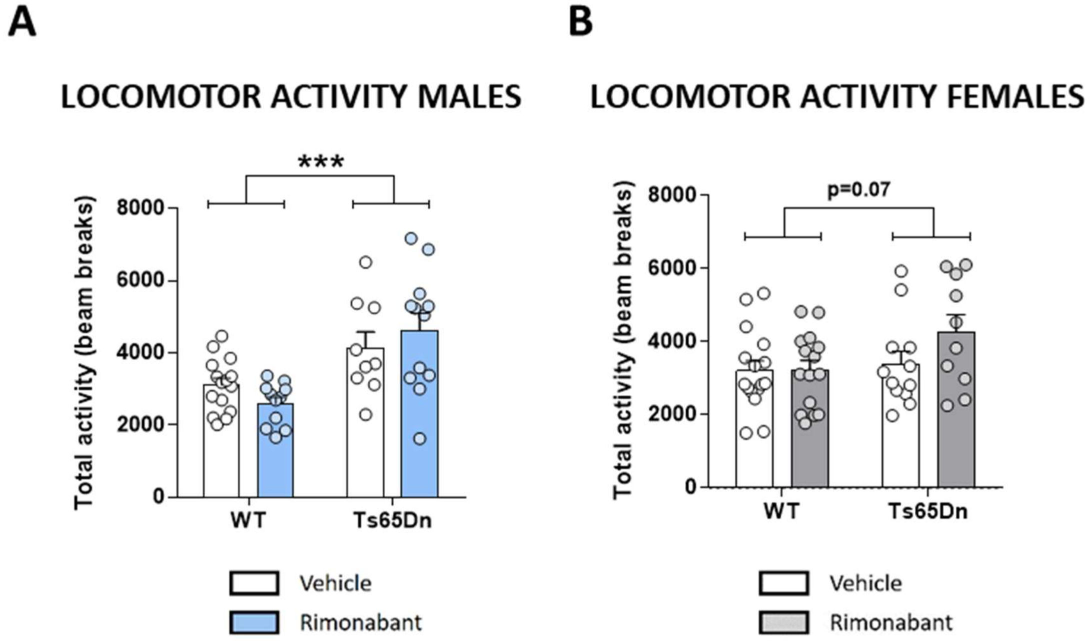


**Supplementary Figure 2. Long-term CB1R pharmacological inhibition does not modify locomotor activity in Ts65Dn.** (A-B) Total activity measured as number of beam breaks of male (A) and female (B) WT and Ts65Dn mice of 5 months of age treated with VEH or RIM (0.5 mg/kg/day) (males: WT VEH, n=15; WT RIM, n=11; Ts65Dn VEH n=9; Ts65Dn RIM n=12; females: WT VEH, n=16; WT RIM, n=15; Ts65Dn VEH n=12; Ts65Dn RIM n=10). Distribution of individual data with mean ± s.e.m. *** $p < .001$ (genotype effect) by two-way ANOVA.
